# “The green peach aphid, *Myzus persicae,* transcriptome in response to a circulative, nonpropagative polerovirus, *Potato leafroll virus*”

**DOI:** 10.1101/2021.04.15.440077

**Authors:** MacKenzie F. Patton, Allison K. Hansen, Clare L. Casteel

**Author notes:** Corresponding author: Clare L. Casteel, Email Address (CLC).

## Abstract

Viruses in the *Luteoviridae* family, such as *Potato leafroll virus* (PLRV), are transmitted by aphids in a circulative and nonpropagative mode. This means the virions enter the aphid body through the gut when they feed from infected plants and then the virions circulate through the hemolymph to enter the salivary glands before being released into the saliva. Although these viruses do not replicate in their insect vectors, previous studies have demonstrated viruliferous aphid behavior is altered and the obligate symbiont of aphids, *Buchnera aphidocola,* may be involved in transmission. Here we provide the transcriptome of green peach aphids (*Myzus persicae*) carrying *PLRV* and virus-free control aphids using Illumina sequencing. Over 150 million paired-end reads were obtained through Illumina sequencing, with an average of 19 million reads per library. The comparative analysis identified 134 differentially expressed genes (DEGs) between the *M. persicae* transcriptomes, including 64 and 70 genes that were down- and up-regulated in aphids carrying PLRV, respectively. Using functional classification in the GO databases, 80 of the DEGs were assigned to 391 functional subcategories at category level 2. The most highly up-regulated genes in aphids carrying PLRV were cytochrome p450s, genes related to cuticle production, and genes related to development, while genes related to histone and histone modification were the most down-regulated. PLRV aphids had reduced *Buchnera* titer and lower abundance of several *Buchnera* transcripts related to stress responses and metabolism. These results suggest carrying PLRV may reduce both aphid and *Buchnera* genes in response to stress. This work provides valuable basis for further investigation into the complicated mechanisms of circulative and nonpropogative transmission.

## Introduction

Aphids are a member of the superfamily Aphidoidea, are distributed world-wide, and cause major damage to global agricultural (1). Despite there being over 4,000 species, only about 400 are known as significant pests (2). Aphids are effective pests partially because they do not require sexual reproduction and can use parthenogenesis to quickly increase their numbers (3). Another aspect of aphid biology that makes them an effective pest is their host range. Although many aphids are very specialized herbivores, only feeding on a few related species, some species feed on many taxa of plants (4). *Myzus persicae* is one of these polyphagous pests, feeding on over 40 different families, including Solanaceae (5).

Along with causing direct feeding damage, aphids are important plant virus vectors, representing over 50% of all known insect vectors for plant viruses (6)(7,8). Increasing evidence has shown that plant viruses alter vector host finding, dispersal, and inoculation through changes in host physiology, however the underlying mechanisms are largely unknown (9,10)(11). Recent evidence suggests that viruses may also directly affect aphid biology (12–15). *Rhopalosiphum padi* aphids carrying *Barley yellow dwarf virus* (BYDV), a virus from the genus *Luteovirus*, prefer non-infected hosts, while virus free *R. padi* prefer infected hosts (12). In these experiments, the authors determined this interaction was due to direct impacts on the aphid and not due to virus induced changes in host plant quality by purifying BYDV virions and feeding them to aphids in artificial diet before conducting aphid bioassays.

Although their lifestyle or host may change, all aphids depend on *Buchnera aphidicola* as their primary obligate endosymbiont (16). *Buchnera* provide the aphid with essential amino acids and nutrients that are limited in the aphid’s diet (17–20), and because of this aphids can no longer survive without *Buchnera.* For example, when *Buchnera* is reduced by using antibiotics, studies have shown lower body mass, lower fecundity, and changes to feeding behavior (21,22). Essential components of the *Buchnera* genome have been lost (23,24) as *Buchnera* co-diversified with aphids over time (16,25,26). Because of this *Buchnera* also depends on aphids for survival, living inside special aphid cells, known as “bacteriocytes”. Together, however, this mutualistic duo wreaks havoc on agriculture across the globe.

Previous studies have speculated *Buchnera* may have a role in aphid transmission of plant viruses (26–30). Specifically, the *Buchnera* chaperone protein GroEL, a homologue from *Escherichia coli* (31), has been implicated in transmission for a number of viruses (26,27,30,32–35). Direct interactions are thought to be unlikely due to the spatial separation of bacteriocytes and circulating virions (26). However, GroEL from *Buchnera* is found in aphid saliva and has been shown to trigger plant defenses and reduce aphid fecundity using transgenic plants expressing GroEL (36,37). Virus-induced changes in plant defense or nutrients may alter aphid-*Buchnera*-plant interactions, highlighting the need for additional research on virus-vector-*Buchnera* interactions (38).

*Potato leafroll virus* (PLRV) is a positive sense ssRNA virus and the type member of the genus *Polerovirus* (family *Luteroviridae*). PLRV is phloem limited and transmitted in a circulative nonpropagative form. This means the virus particles will travel across the gut membrane on specific receptors into the insect hemolymph. From here it will traverse to the salivary gland and duct so that it may be injected back into the phloem tissue (9,39). Previous studies have shown that aphid vectors prefer to settle on plants infected with PLRV and that insect vectors have higher fecundity when feeding on these plants compared to controls (5,40–42). Additionally, a recent study investigating aphid small RNAs (sRNAs), found that *M. persicae* who fed upon PLRV infected plants and purified PLRV diets had significantly altered sRNA profiles, manipulated immune response, and differentially expressed *Buchnera* sRNAs that are associated with tRNAs (43). The role of *Buchnera* tRNA associated sRNAs is largely unclear, as their expression is conserved in divergent *Buchnera* taxa (44), they are differentially expressed during aphid development (45) and when aphids feed on different host plant diets (46). In consequence, the mechanisms mediating virus-aphid-*Buchnera* interactions are largely unknown. Recently we demonstrated PLRV induces changes in plant nutrients and defenses in infected host plant, (47) however, the impacts of these changes on symbiont-aphid interactions are unknown. To address this lack of knowledge we examined changes in the transcriptome of *M. persicae* with and without PLRV, *Buchnera* titer, and changes in aphid and *Buchnera* transcripts from aphids feeding on PLRV-infected plants. By providing evidence that nonpropagative circulative plant viruses can affect insect vectors through changes in the transcriptome and alter *Buchnera* titer, our study will contribute to growing knowledge of the insect microbiome at a plant-insect interface.

## Methods

### Plant and insect growth conditions

*Solanum tuberosum* were propagated using leaf-bud cutting from cv. Désirée (48) in laboratory experiments. Plants were grown in growth chambers under controlled conditions (25/23°C day/night with a photoperiod of 16/8 h day/night). Non-viruliferous and viruliferous aphid clones of a potato-adapted red strain of *Myzus persicae* were reared under controlled conditions (25/23°C day/night with a photoperiod of 16/8 h day/night) on healthy potato. All experiments were conducted in the same environmental chambers and conditions, so there were no environmental differences in treatments (25/23°C day/night with a photoperiod of 16/8 h day/night).

### Pathogen infection

*Agrobacterium tumefaciens* (LBA4404) containing the infectious clone of PLRV (49) was grown at 28°C in in LB broth (+10 mM MgSO4), with kanamycin (50 μg/mL), carbenicillin (100 μg/mL) and rifampicin (50 μg/mL) for selection. After 24 hours, bacteria were centrifuged to concentrate and resuspended in 10mM MgCl2. One week old *S. tuberosum* were inoculated at an optical density (OD) of 0.70. Three weeks post infection, tissue was collected from all plants, RNA was extracted using the SV Total Isolation Kit as per manufacturer’s instructions (Promega, Madison, WI, USA), and cDNA was synthesized using 1500 ng of total RNA and random hexamers (20ng/μL) with the SMART® MMLV as per manufacturer’s instructions (Takara Bio USA, Mountain view, CA, USA). cDNAs were used in PCRs with PLRV specific primers (F-5’ATGAGTACGGTCGTGGTT-3’ and R-5‘CTATTTGGGGTTTTGCAAAGC-3’). A set of uninfected potato cuttings were grown at the same time as the plants above to serve as controls. After systemic plant infection was verified plants were immediately used in experiments.

### RNAseq, qRT-PCR, and qPCR aphid experiments

Five adult aphids were placed on the first fully expanded leaflet of three infected and on three healthy plant. After 24 hours, all adults were removed and 20 larvae were left to develop. Seven days later all aphids were at the same developmental stage and 10 young aphids were collected into a tube from each plant (N=3 plants with 10 aphids per plant) and immediately frozen in liquid nitrogen until use in RNAseq experiments. The entire experiment was repeated a second time for confirmation of RNAseq results using qRT-PCR and to examine the titer of the bacterial symbiont, *Buchnera*, in aphids. For this experiment 5 aphids were collected for RNA extraction and 5 aphids were collected for DNA extractions from each plant. We also prepared 6-7 plants for each treatment instead of 3 (N = 6-7 with 5 aphids per plant), however all other methods were the same.

### RNA and DNA isolation from aphids

RNA was extracted from aphid tissue collected in the first two experiments as described above. The RNA concentration and purity were measured using a NanoDrop. The integrity of RNA was confirmed using the Bioanalyzer 2100 system (Agilent, Santa Clara, CA, USA). DNA was extracted from aphid tissue collected in the second experiment using Cetyl trimethylammonium bromide. The integrity of DNA was confirmed using an agarose gel. The DNA concentration and purity were measured using a NanoDrop (Thermo Fisher Scientific, Waltham, MA, USA).

### Library Preparation, and Sequencing

Sequencing libraries were prepared using a multiplexing library protocol (50). Briefly, oligo(dT) 25 Dynabeads were used to purify mRNA, which was then fragmented, and the first-strand cDNA was synthesized using random primers, dNTP, and reverse transcriptase. The second-strand was synthesized using a dUTP mix, DNA Polymerase I, and RNase H, ends repaired, and adenylated. The cDNA fragments were ligated to adapters, selectively enriched by PCR, and purified using the AMPure XP beads. The library quality was assessed using the Agilent Bioanalyzer 2100 system and sequenced using an Illumina HiSeq 2000 instrument.

### Read mapping, differential expressed gene (DEG) analysis, and Gene Ontology (GO) classification

RNA-Seq data were analyzed using RStudio (Version 1.1.383) and Bioconductor according to Anders et al. (2013) with some modifications (48) (See Supplementary Figure S1). Sequence quality was determined, trimmed, and poor-quality reads removed using ShortRead (52) and FastQC (53). Reads were mapped to the *Myzus persicae* clone G006 genome v2.0 from AphidBase (54) using TopHat2 (55). Mapped reads were assigned to genes and counted with HTSeq (56) and normalized by size factors obtained from the negative binomial-based DESeq2 package (57). Gene annotation files for both were downloaded from NCBI. After normalization, clusterization profiles of the samples were assessed by hierarchical clustering (with Euclidean distance metric and Ward’s clustering method) and principal component analysis (PCA). Differentially expressed genes (DEGs) between infected and control treatments were identified using DESeq2 (57). Genes with False Discovery Rate (FDR)-corrected p-values ≤ 0.1 were classified as differentially expressed. For Gene ontology (GO) analysis, Blast2GO software (58) was utilized for annotation as previously described (59).

### qRT-PCR

cDNA was synthesized using random hexamer (20ng/μL) and quantitative RT-PCR (qRT-PCR) was performed. Transcript abundance was quantified for the *M. persicae* genes, *HSP68-like* (MYZPE13164_G006_v1.0_000070430.1) and *M. persicae* cuticle protein5-like (*MPCP5-like)* (MYZPE13164_G006_v1.0_000133030.2), and for the *Buchnera* genes, *argE (BUMPUSDA_CDS00542), dnaK (BUMPUSDA_CDS00441),* and *groEL (BUMPUSDA_CDS00567),* using gene specific primers (Supplementary Table S1). qRT-PCR reactions were carried out using SYBR green PCR master mix (Applied Biosystems, Carlsbad, CA, USA), in an CFX384 instrument (Bio-Rad, Hercules, CA, USA). Three technical replicates were performed for each individual sample, and a digital pipette was used for all pipetting. Relative transcript abundance was calculated utilizing a standard curve produced from 10-fold series dilution of cDNA synthesized from 1000 ng/μL of total RNA according to the standard curve method (Applied Biosystems, Carlsbad, CA, USA). Technical replicates of raw CT values were averaged and transcripts of interest were normalized to the house-keeping transcript ribosomal protein L7 *rpl7* for aphids or the 50S ribosomal subunit gene *rpIN* for *Buchnera*, as previously described (60–62).

### *Buchnera* titer

*Buchnera* titer here is defined as the ratio of *Buchnera* single copy genes to aphid single copy genes. To determine *Buchnera* titer in whole aphid bodies we used quantitative PCR (qPCR) and measured the ratio of a single copy *Buchnera* gene (*rplN*) to a single copy aphid gene (*RPL7*). qPCR reactions were carried out using SYBR green PCR master mix (Applied Biosystems, Carlsbad, CA, USA), in the CFX384 instrument (Bio-Rad, Hercules, CA, USA). Reactions were performed in triplicate for each sample, and the average was used for analysis. Relative abundance was calculated utilizing a standard curve produced from 10-fold serial dilution of DNA.

### Statistical analyses

RNAseq data analyses were performed as described above. All statistical analyses for qRT-PCR and qPCR were determined using a *P* < 0.05. Analysis of variance (ANOVA) was used to determine significant difference in transcript abundance. For *Buchnera* titer single factor ANOVA was used to determine difference in relative abundance. The statistical analyses were performed using JMP 8 software (SAS Institute, Cary, NC, USA).

## Results

### Differential gene expression in the presence of PLRV

Over 150 million paired-end reads were obtained through Illumina sequencing, with an average of 125,815, 247 100-bp reads per library (Table 1, Fig 1A). About 73% of the reads mapped to the *M. persicae* reference genome, with around only 27% being uniquely mapped (Table 1, Fig 1B). In order to examine biological variability, a principal component analysis (PCA) of the normalized count data was performed (Fig 1C). The first component of variance separated samples by treatments and accounted for 54% of the variance. Hierarchical clustering confirmed PCA results in visual representation of DEG expression (Fig 1D). The transcriptome of viruliferous *M. persicae* was compared to the transcriptome of virus-free *M. persicae* using the negative binomial-based DESeq2 (57). Overall, 96 differentially expressed genes (DEGs) were detected using an FDR adjusted p-value ≤ 0.05 and log2 fold change (FC) ≥ 0.5, however, by relaxing our FDR adjusted p-value to ≤ 0.1 we were able to include 38 additional DEGs (134 DEGs total included; Fig 1E; Supplementary Table S2). The presence of PLRV in the aphid vector caused a down-regulation of 70 genes and an up-regulation of 64 genes (Fig 1F; FDR adjusted p-value ≤ 0.1 and log2 fold change (FC) ≥ 0.5). Overall, 0.8 % of the aphid genome was significantly impacted by the presence of PLRV.

**Table 1:** RNAseq Stats.

| Sample Name | Total paired-end reads | Total alignments | Aligned | Unique paired | Non-unique paired |
| --- | --- | --- | --- | --- | --- |
| Control 33 | 40,676,571 | 30,990,077 | 76.17% | 28.52% | 47.67% |
| Control 34 | 18,942,570 | 13,440,120 | 70.80% | 25.89% | 45.07% |
| Control 35 | 22,651,445 | 16,478,122 | 72.49% | 25.93% | 46.82% |
| PLRV 36 | 25,343,473 | 18,306,992 | 72.12% | 26.30% | 45.94% |
| PLRV 37 | 23,743,718 | 17,496,940 | 73.63% | 27.15% | 46.54% |
| PLRV 38 | 23,533,704 | 17,410,337 | 73.78% | 27.05% | 46.93% |

**Fig 1.**
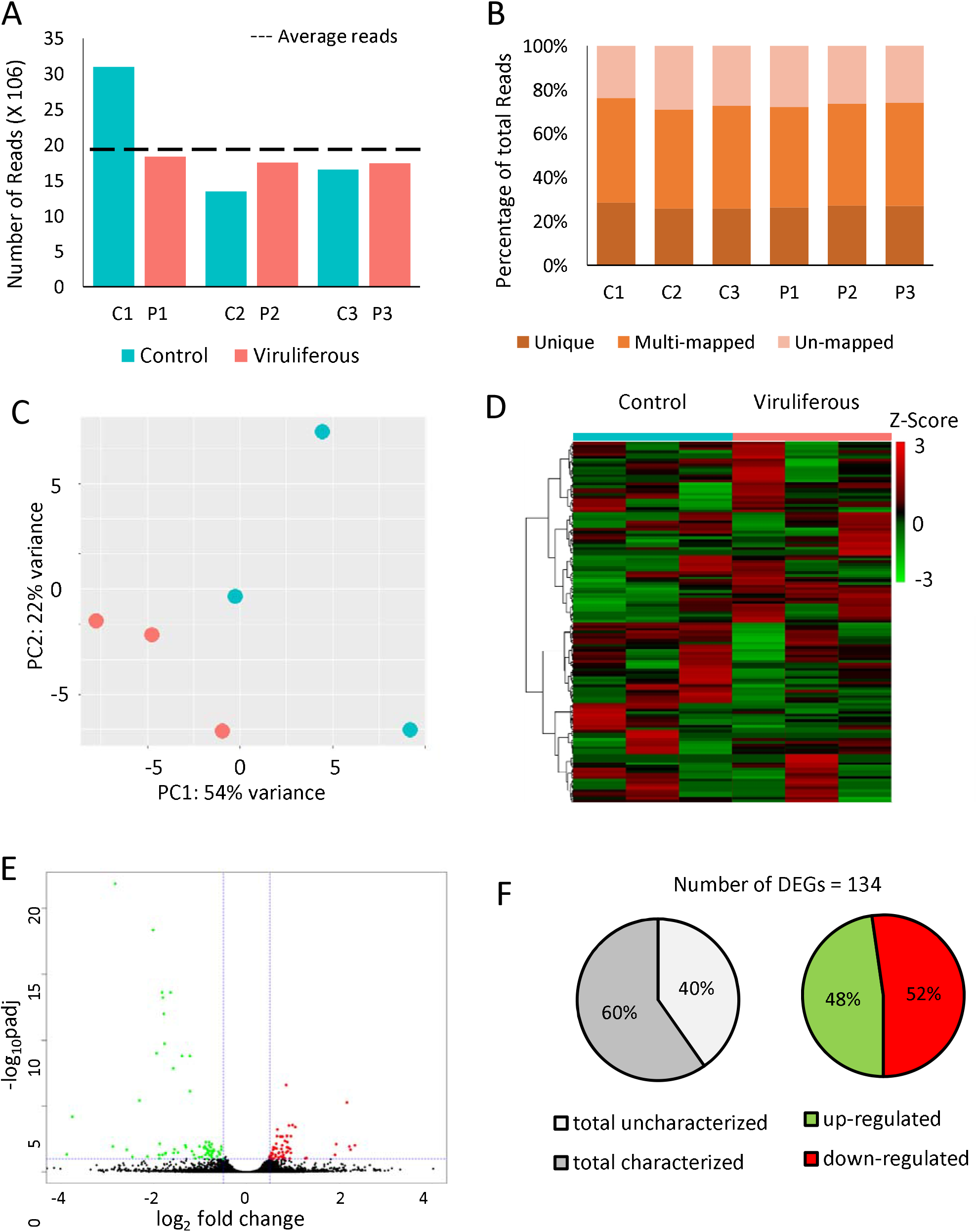
Overview of *Myzus persicae* transcriptome after *Potato leafroll virus* (PLRV) acquisition. (A) Number of paired-end reads generated for each library by Illumina HiSeq sequencing. The dashed line represents the average of paired-end reads from all 6 libraries. (B) Proportion of uniquely mapped, multimapped, and unmapped reads obtained for each library. Reads were mapped in the *Myzus persicae* clone G006 genome (AphidBase). (C) Principal component analysis of normalized count data from all samples. (D) Hierarchical clustering analysis of normalized count data z-scores exhibited by differentially expressed genes (DEGs) within each sample. (E) Volcano-plots of −log10p and log2FC exhibited by each gene in viruliferous aphids compared to controls. Up- and down-regulated genes are presented in red and green, respectively. (F) Numbers of up- and down-regulated DEGs in viruliferous aphids in comparison to control aphids. DEGs were identified using DESeq2 and defined by |log2FC| ≥ 0.5 and false discovery rate (FDR)-corrected p-value ≤ 0.1. Control (aphids without virus); Viruliferous (aphids carrying PLVR). FC, fold-change; p, FDR-corrected p-value.

### Functional roles of differentially expressed genes

Gene ontology (GO) enrichment analyses were performed with the DEGs from each treatment to identify functions and pathways disturbed in aphids carrying PLRV. One or more gene ontology terms were assigned to each transcript from biological processes, molecular functions, and cellular compartments term using Blast2GO functional gene annotation (Conesa et al. 2005). The 134 DEGs were assigned to functional GO terms within the three categories, including 125 biological processes, 118 molecular functions, and 148 cellular compartments. Of the 134 DEGs, 53 (39.55% of total DEGs) were classified as “uncharacterized proteins.” The majority of DEGs assigned to biological processes were categorized as metabolic processes (41%), cellular-protein processes (11%), and oxidation-reduction processes (11%) (Fig 2A). As for DEGs assigned to molecular functions, almost half were associated with catalytic activity (33%) or nucleic acid binding (21%) (Fig 2B). Within the cellular component category, 24% were related to the membrane and 24% were related to intracellular locations (Fig 2C).

**Fig 2.**
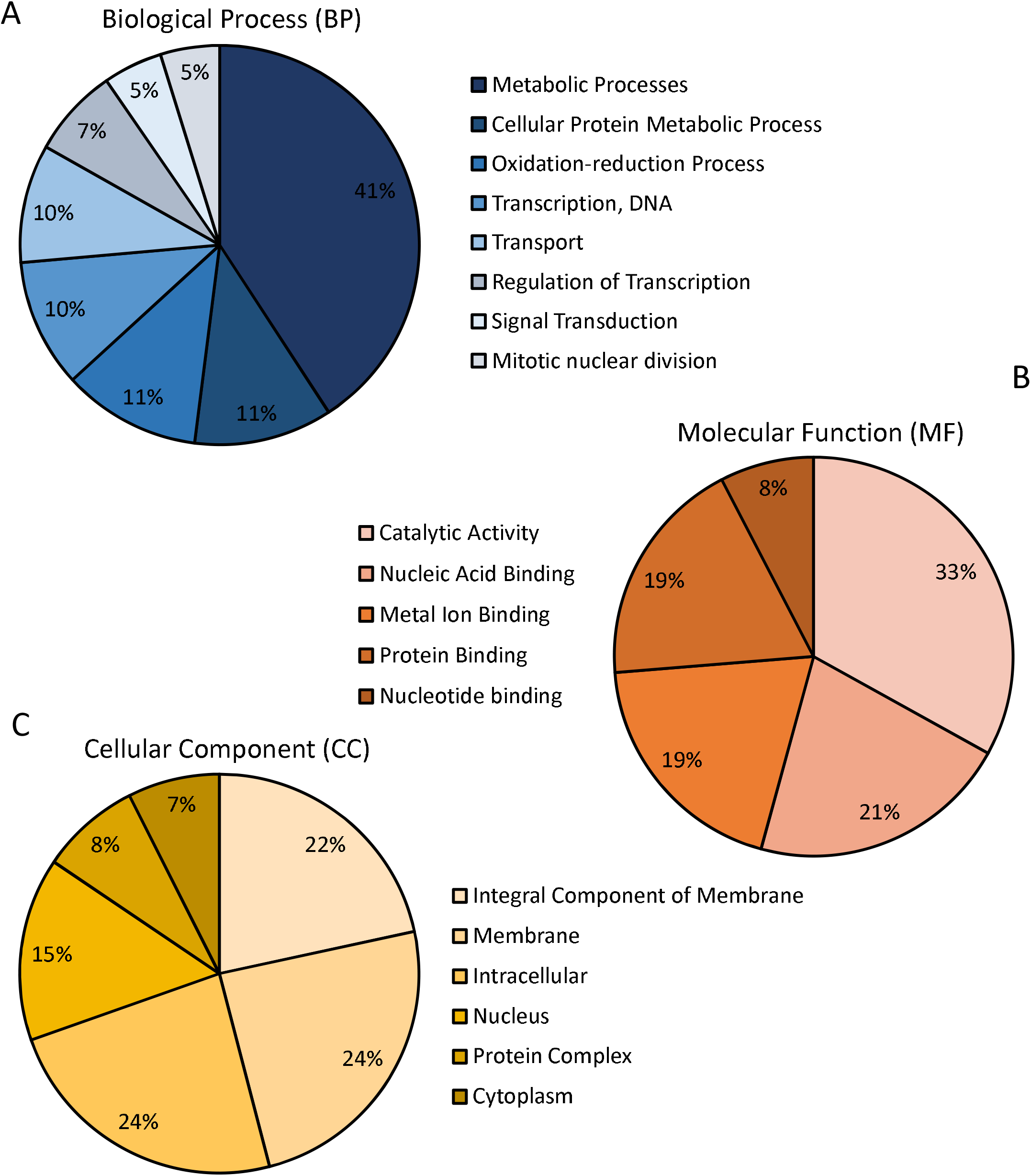
Blast2GO Gene Ontology of DEGs arranged by functional categories. (A) Biological processes (BP), (B) Molecular Function (MF), and (C) Cellular Component (CC). The predicted gene functions of differentially expressed genes as assigned by Blast2GO at level 2-3 in each aforementioned category. Each DEG may be assigned to one or more GOterm, with a total of 391 GOterms from the three functional groups assigned to the 134 DEGs.

Next each DEGs was annotated using a single Blast2GO consensus description. Many of the genes up-regulated in PLRV aphids were related to cuticle formation and development (16%) and catalytic activity (16%), however the majority of up-regulated transcripts were uncharacterized (31%; Fig 3A). The largest groups of down-regulated genes in PLRV aphids were related to histones (10%), catalytic activity (10%), transmembrane transport (9%), proteolysis or protein ubiquitination (7%), and nucleic acid binding and metabolic processes (7%; Fig 3B). A significant proportion of the down-regulated transcripts in PLRV aphids were also uncategorized (47%). The most highly expressed DEGs included transcripts related to cuticle formation and development and 4C1-like cytochrome P450s (Table 2). The most down-regulated transcripts in PLRV aphid were related to histones and histone modifying proteins (Table 3).

**Table 2:**
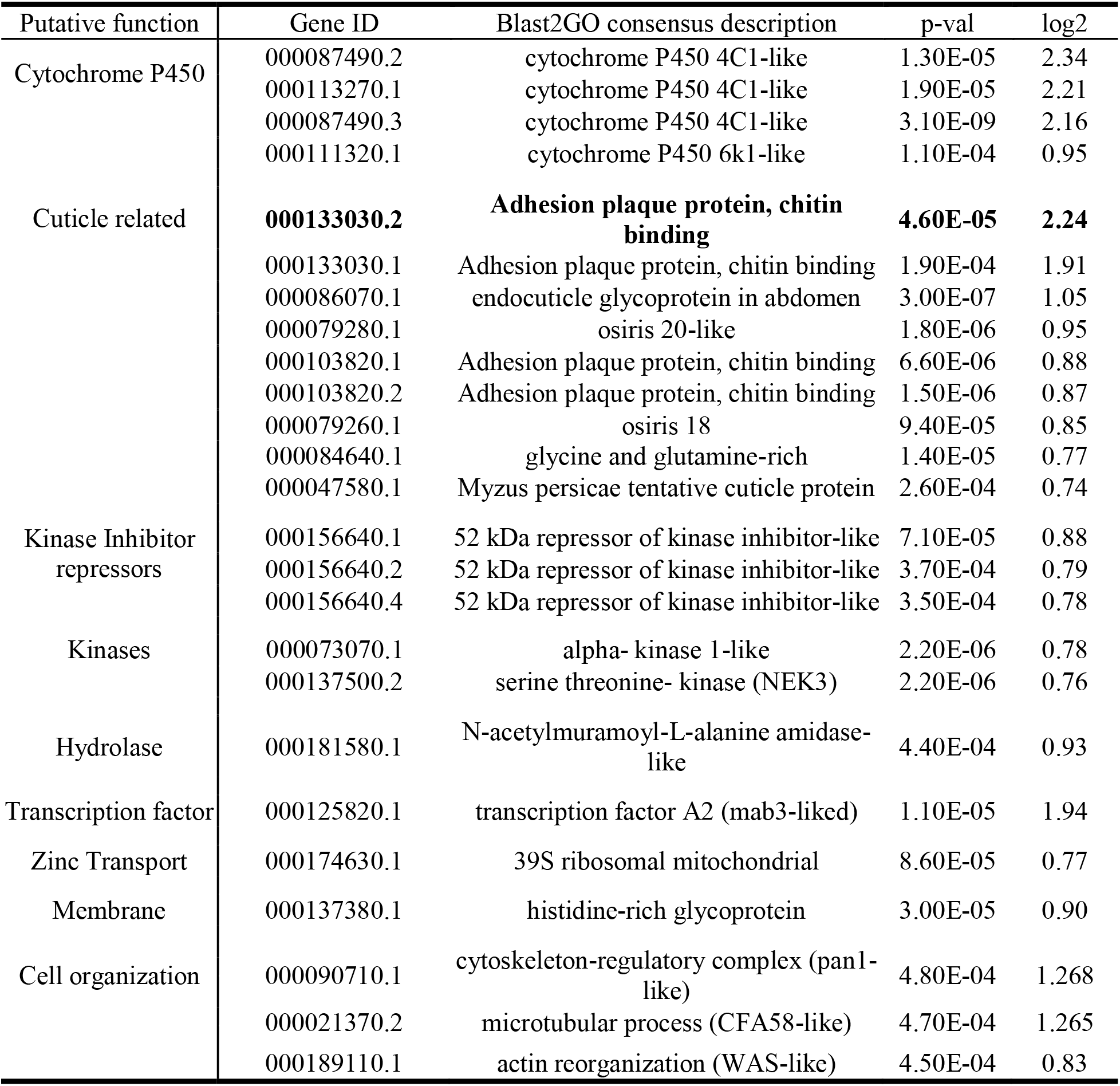
Most highly up-regulated *M. persicae* genes that were characterized* in aphids carrying PLRV compared to controls. DEGs determined by adjusted p value < 0.1 and described by Blast2GO. Gene ID corresponds to MYZPE13164_G006_v1.0_XXXXXXXXX.X found on AphidBase.org. Regulation of bolded transcripts were validated in a separate experiment. *20 uncharacterized genes were up-regulated (See Supplementary Table S2)

**Table 3:** Most highly down-regulated *M. persicae* transcripts that were characterized* in aphids carrying PLRV compared to controls. DEGs determined by adjusted p value < 0.1 and described by Blast2GO. Gene ID corresponds to MYZPE13164_G006_v1.0_XXXXXXXXX.X found on AphidBase.org. Regulation of bolded transcripts were validated in a separate experiment. *33 uncharacterized genes were down-regulated (See Supplemental Table S2)

| Putative function | Gene | Blast2GO consensus description | p-value | log2 |
| --- | --- | --- | --- | --- |
| Histones | 000100490.1 | histone H3 | 5.06E-05 | -2.58 |
|  | 000100610.1 | histone H3 | 2.69E-04 | -2.14 |
|  | 000100770.1 | histone H3 | 1.04E-04 | -1.74 |
|  | 000100600.1 | histone H2A | 1.38E-04 | -1.75 |
|  | 000092680.1 | histone H2A-like | 9.76E-05 | -1.47 |
|  | 000100590.1 | histone H2B-like | 7.37E-05 | -1.18 |
|  | 000100620.1 | histone H4 | 3.02E-04 | -0.96 |
| Histone Modifying | 000163990.2 | glycine-rich DOT1-like | 7.33E-05 | -0.81 |
|  | 000163990.1 | glycine-rich DOT1-like | 1.97E-04 | -0.76 |
| ubiquitination | 000119640.3 | E3-ubiquitin ligase RNF19B-like | 6.08E-06 | -0.88 |
|  | 000119640.1 | E3-ubiquitin ligase RNF19B-like | 1.42E-05 | -0.81 |
|  | 000119640.2 | E3-ubiquitin ligase RNF19B-like | 2.34E-05 | -0.78 |
| Hydrolase | 000133360.1 | serine carboxypeptidase | 1.10E-04 | -1.58 |
|  | 000083200.2 | Arylsulfatase B-like | 4.69E-04 | -1.11 |
|  | 000083200.1 | Arylsulfatase B-like | 3.25E-04 | -1.02 |
|  | 000200070.1 | Thioesterase (THEM6-like) | 1.13E-04 | -0.82 |
| Response / Immunity | 000071560.1 | Protease inhibitor (Papain inhibitor) | 2.91E-04 | -2.46 |
|  | **000070430.1** | **Heat shock 68-like** | **4.85E-04** | **-1.88** |
|  | 000193260.2 | G- coupled receptor Mth-like 3 | 2.03E-04 | -0.80 |
| Transport | 000029670.1 | dynein intermediate chain-like | 1.90E-05 | -1.01 |
|  | NRF6 | Lipid transport (NRF6-like) | 5.18E-05 | -0.85 |
|  | 000203490.1 | zinc finger C3H1 type-like 2-A | 1.03E-04 | -0.86 |
|  | 000072950.2 | Sugar transport (TRET1-like) | 6.74E-06 | -0.81 |
| nucleic acid metabolism | 000036830.1 | DNA integration<br>(pol poly retrotransposon-related) | 1.86E-04 | -0.89 |
|  | 000012610.1 | mRNA catabolic process<br>(BRISC/BRCA1-A complex-Like) | 5.14E-04 | -0.84 |

**Fig 3.**
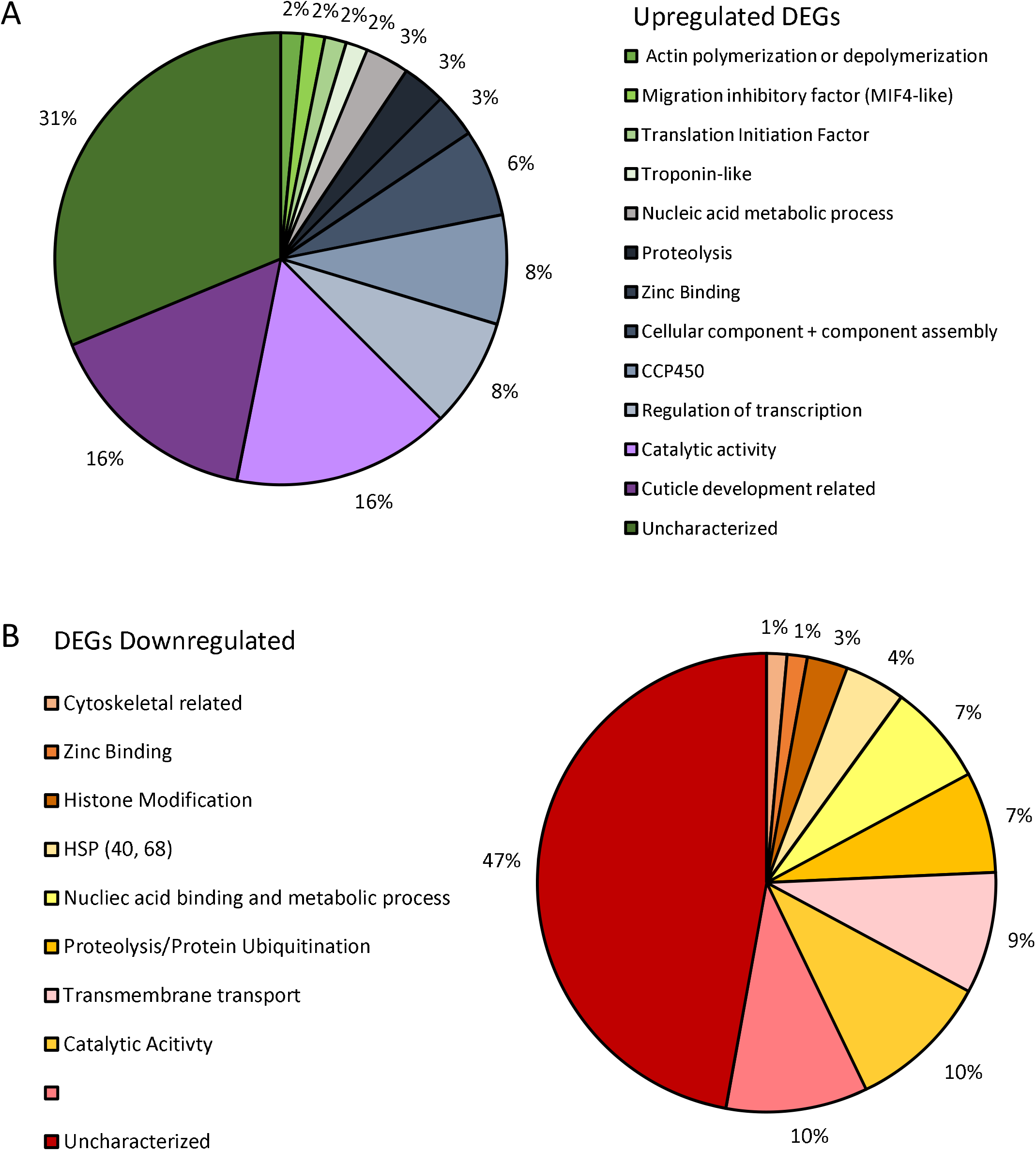
Blast2GO annotation for up-regulated DEGs and down-regulated DEGs in aphids carrying PLRV compared to controls. The consensus description predicted by Blast2GO of the (A) 64 up-regulated DEGs and the (B) 70 down-regulated DEGs.

### Validation of aphid transcripts via RT-qPCR

To validate the RNAseq analysis a separate experiment was performed using the same experimental design and the transcript abundance of one up-regulated gene (MYZPE13164_G006_v1.0_ 000133030.2 (*MPCP5-like*)) and one down-regulated gene (MYZPE13164_G006_v1.0_ 000070430.1 (*HSP68-like*)) was measured using qRT-PCR (bolded genes in Table 2 and 3). Consistent with the RNA-seq data, abundance of the *MPCP5-like* transcript was significantly higher in viruliferous *M. persicae* compared to virus-free controls (10.045, 2.017, relative expression respectively; p = 0.019; Fig. 4A). Abundance of the *HSP68-like* transcript was significantly lower in viruliferous *M. persicae* when compared to virus-free controls (1.44, 5.51, relative expression respectively; p < 0.01) (Fig 4A-B).

**Fig 4:**
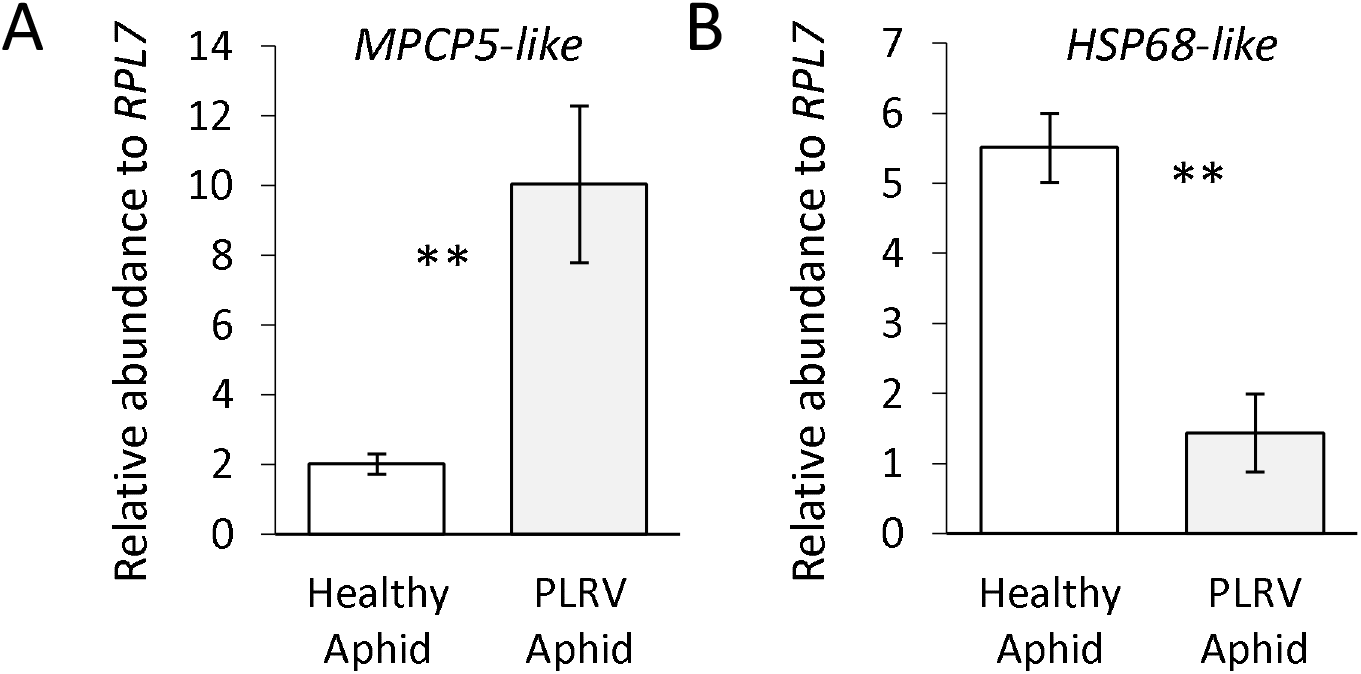
Relative transcript abundance of two genes in *Myzus persicae* with and without *Potato leafroll virus* (PLRV). (A) A cuticle related protein (MPCP5-like) was significantly up-regulated in expression in individuals with PLRV compared to virus free controls. (B) A predicted heat shock protein (HSP68-like) was significantly down-regulated in expression in individuals with PLRV compared to controls. Transcripts were measured relative to a housekeeping gene *RPL7*. Significant differences were calculated using an ANOVA (\**P* < 0.05; Error bars represent ±SEM).

### The impact of PLRV on *Buchnera aphidicola* titer

*Buchnera* has been previously implicated in transmission of PLRV and other luteoviruses (26,28,30,34,63), however *Buchnera* titer and coding sequence transcripts have not been examined in aphids carrying PLRV. From our experiments, *Buchnera* titer was ~1.5 times higher for virus-free aphids compared to aphids carrying PLRV (ratios 6.42, 4.20 respectively; p = 0.037; Fig 5A). To investigate the potential mechanisms mediating decreases in *Buchnera* titer we measured abundance of two transcripts related to stress, *dnaK* (64,65) and *groEL* (65,66), and one transcript related to metabolism, *argE* (45). Abundance of all three transcripts were reduced in aphids carrying PLRV compared to controls. Viruliferous aphids had 36.68% less *argE* transcripts (p = 0.026), 16.67% less *groEL* transcripts (p = 0.024), and 18.77% less *dnaK* transcripts (p = 0.046) compared to that of the virus free aphids (Fig 5B-D).

**Fig 5.**
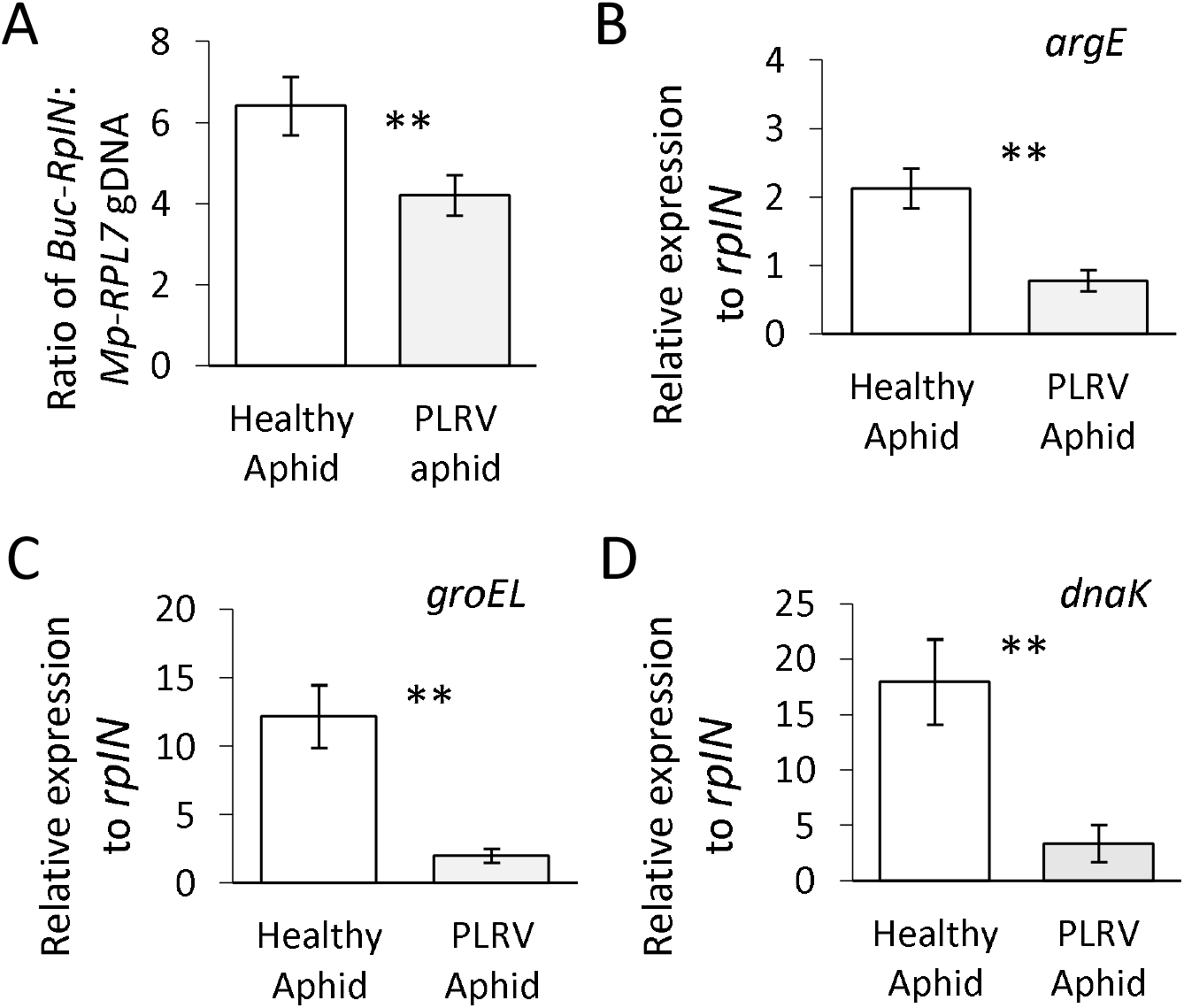
*Buchnera aphidicola* titer and transcript changes in *Myzus persicae* with *Potato leafroll virus* (PLRV). (A) Ratio of a single copy *Buchnera* gene *rpIN* relative to a single copy aphid gene *RPL7,* demonstrates *M. persicae* with PLRV have a decreased *Buchnera aphidicola* titer relative to control aphids. (B-D) *Buchnera* transcripts *groEL*, *dnaK*, and *argE* relative to the *Buchnera* gene *rpIN* housekeeping gene. All three transcripts were down-regulated in viruliferous *M. persicae* compared to virus free controls. Significant differences were calculated using an ANOVA (\**P* < 0.05; Error bars represent ±SEM).

## Discussion

The main focus of this paper was to examine the effect that *Potato leafroll virus* has on the transcriptome of *M. persicae,* and their primary endosymbiont *Buchnera aphidicola*. The largest category of known up-regulated transcripts in viruliferous aphids compared to controls were related to the cuticle and cuticle development. Insect cuticles are largely composed of a protein matrix embedded with chitin filaments (67). Cuticle proteins (CPs) have been shown to be involved in general development, molting, transmission of non-persistent viruses, and insecticide resistance through changes cuticle permeability (33,68–71). In *Acyrthosiphon pisum*, 19 CPs were found to be regulated by photoperiodism and suspected to be involved in the transition from asexual to sexual production (72). Further, cuticle proteins have been implicated as potentially facilitating transmission of *Barley yellow dwarf virus* (BYDV-GPV), *Cereal yellow dwarf virus* (CYDV-RPV), and *Turnip yellows virus* (TuYV), three related Luteoviridae viruses (73–75). Whilst we cannot know the function of changes in CP transcripts in PLRV-aphid interactions from these experiments, these genes represent promising targets for further investigation.

In addition to many cuticle related proteins, five *cytochrome P450s* genes were significantly up-regulated in viruliferous aphids compared to controls. Cytochrome P450s play important roles in hormones and pheromones metabolism but are more famous for their roles in the metabolism of insecticides and host plant chemicals. Polyphagous insects, like *M. persicae,* encounter many different hosts and tend to have high numbers P450-based metabolism of allelochemicals compared to more specialized aphids (76). Previous work has shown that a *cytochrome 450 gene* (*CYP6CY3*) was found to increase nicotine tolerance and aphid host adaptation (77,78). It has been previously hypothesized that upregulation of p450s could help insect vectors tolerate less desirable hosts which could be beneficial to the virus (79).

Transcripts encoding a heat shock protein (HSP68-like) was among the most down-regulated in virluiferous aphids compared to controls. HSP68 is a member of the HSP70 family, which are important chaperone proteins that are known to be up-regulated in response to stress. One study found that the *HSP70* from *Bemisia tabaci* is up-regulated after acquisition of *Tomato yellow leaf curl virus* (TYLCV) (80). They went on to show that HSP70 protein can directly interact with TYLCV using *in vitro* studies and that they co-localize together in insect midgut cells using *in situ* hybridization. The authors suggest HSP70 may play an inhibitory role in virus transmission, as transmission was reduced when whiteflies were fed HSP70 antibodies. Because *HSP68-like* transcripts were down-regulated in aphids carrying PLRV in our study, it’s tempting to speculate that this may increase PLRV transmission. Porras et al. demonstrated that BYDV-PAV, a strain that is only transmitted by *Rhopalosiphum padi* (bird-cherry oat aphid), up-regulated the abundance of three *HSP70* transcripts in the aphid vector. The authors found BYDV infection increases plant surface temperature and aphid heat tolerance, suggesting a protective role of HSP70 proteins in virus-aphid-plant interactions (81). Although it is not known if PLRV increases plant surface temperature and vector heat tolerance, it has been shown that potato plants kept at higher temperatures are more susceptible to PLRV than compared to lower temperatures (82). Also aphid acquisition and transmission at higher temperatures has previous resulted in higher transmission rates compared to lower temperatures (83), however at very high temperatures differences were reduced (84). It is not known how decreases in *HSP68-like* transcripts in aphids carrying PLRV in our study may alter aphid heat tolerance.

Several histone genes were down-regulated in viruliferous *M. persicae* compared to controls in this study. Histones are involved in DNA organization and regulation of gene expression (85), but have also been shown to be induced in response to stress and starvation (86). Histone depletion can lead to changes to an open chromatin configuration and large scale shifts in expression, however it should be noted histone depletion is also associated with DNA damage (87). In a recent study investigating small RNAs (sRNAs), *M. persicae* who fed upon PLRV infected plants and purified PLRV diets had significantly altered sRNA profiles and immune responses. Ultimately, the significance of this response is unknown, and further investigation is necessary to understand potential impacts of the downregulation of these genes.

In this study there was a significant reduction of *Buchnera* titer and *Buchnera* gene expression of three genes (*dnaK*, *groEL,* and *argE*) in aphids carrying PLRV compared to control aphids. In general, gene regulation at the mRNA level in *Buchnera* is thought to be minimal because *Buchnera* transcription factors are reduced (88) and very few transcriptional responses had been observed previously (17). Only two transcription initiation factors (σ32 and σ70), the heat shock and housekeeping transcription factors, respectively, remain in *Buchnera* Myzus’s genomes (89) similar to other *Buchnera* taxa (90). The housekeeping sigma factor (σ70) initiates transcription of *argE* which is regulated by the repressor *argR* when bound to arginine in *Escherichia coli* (91). Similar to other Buchnera taxa *Buchnera* Myzus’s genomes (89) has lost the repressor argR so it is unclear how this gene is down-regulated in virus infected aphids compared to un-infected aphids. The other two *Buchnera* genes (*dnaK* and *groEL*) that were down-regulated in this study in aphids carrying PLRV compared to control aphids are associated with the heat shock regulon (90). Moreover, these *Buchnera* genes still retain recognizable σ32 promoter sites up-stream of *dnaK* and *groEL* in the Myzus *Buchnera* G006 genome (NCBI Reference Sequence: NZ_MJNC01000001; Supplemental Table 3) similar to other *Buchnera* taxa (90). The σ32 heat shock response is highly conserved in bacteria and is initiated in response to stress, such as heat shock or other environmental stressors that destabilize proteins (92). In general, compared to free-living bacteria, *Buchnera* only modestly up-regulates genes that still retain the upstream σ32 promoter sites during heat shock (90). In this study it is unclear how PLRV is either directly or indirectly dampening *Buchnera*’s expression of *dnaK* and *groEL* and if it is through a similar mechanism that is also down-regulating the aphid’s stress response genes including *Hsp70*.

A decrease in *Buchnera* titer has previously been associated with different aphid clones (93), plant diets (16), increasing aphid nymphal age (94,95), and heat shock (95–97). For example, a mutation in the promoter region of the heat shock gene (*ibpA*) results in the reduction of *Buchnera* titer (96). Given that the experiments in this study were conducted in the same environment at the same temperature it is highly unlikely that the control aphids experienced heat shock compared to virus infected aphids. Instead, we hypothesize that the virus PLRV is reducing *Buchnera*’s ability to up-regulate genes that are associated with the heat shock regulon and this may lead to increased stress and lysing of *Buchnera* cells and ultimately a reduction of *Buchnera* titer. For example, most obligate pathogens and symbionts, including *Buchnera,* overexpress the protein GroEL during non-heat shock conditions to rescue misfolded proteins (98). Alternatively, as PLRV-infected plants have higher concentrations of free amino acids (47) and we cannot discount the indirect impacts of host plant changes on *Buchnera* titer in our experimental design, similar to Zhang et al. (16) where a change in host plant diet influenced *Buchnera* titer. Buchnera sRNA’s that are hypothesized to regulate *Buchnera* gene expression at the post-transcriptional level have been observed to be differentially expressed when aphids feed on host plants that vary in essential amino acids (46). In turn, aphids carrying PLRV may obtain higher levels of essential amino acids from virus infected plants and as such *Buchnera* genes that are involved in arginine biosynthesis, such as *argE*, are down-regulated compared to control aphids feeding on un-infected plants with lower amounts of essential amino acids.

In other insect-plant pathogen systems plant pathogens are known to modulate obligate symbiont titer. For example, in whiteflies Portiera titer is modulated by the co‐occurrence of its facultative symbiont *Rickettsia* and the Tomato Yellow Leaf Curl Virus (99). In a second system, the obligate symbionts *Carsonella* and *Profftella* of the psyllid *Diaphorina citri* decrease in titer when the plant pathogen *Ca*. Liberibacter asiaticus infects *D. citri* male adults, whereas no significant difference was found in infected female adults (100,101). In adult females’ ovaries, *Carsonella* and *Profftella* titer increased when infected by *Ca.* L. asiaticus. The latter authors hypothesized that endosymbionts titer increases as a part of the psyllid’s immune response and/or response to altered plant nutrition by *Ca.* L. asiaticus infection (100).

In general, this work improves our understanding of the relationships that exist between hosts, viruses, vectors, and endosymbionts, but it also opens up more questions regarding the complexity and depth of these relationships. Aphids and bacterial endosymbionts may benefit from relationships with plant-infecting viruses indirectly or directly but additional studies are needed. Although it is known that *Buchnera* titer and gene expression responses vary with aphid linages (102), it is not known how this is impacted by long term associations with plant-infecting viruses. In regions where virus pressure is high or where poor hosts dominate, aphids may more often be associated with plant infecting viruses. Given the mounting evidence of virus manipulation of insect vectors, this could have lasting impacts on the population structures of these insect vectors and their obligate endosymbiont.

## Acknowledgments

We think Laura Baldwin, Sarah Kate Muehleck, and the many undergraduates that helped with experiments. We thank Gabriella Arena for advice on RNAseq analysis. This work was supported by USDA-NIFA award 2017-67013-26537 and award 2013-2013-03265 to CLC.

## Author contributions

CLC conceived the project. MFP and CLC designed the research. MFP and CLC performed research and analyzed the data. AKH, MFP, and CLC interpreted the data. AKH, MFP, and CLC wrote the article.

## Supporting Information

**Figure S1:**
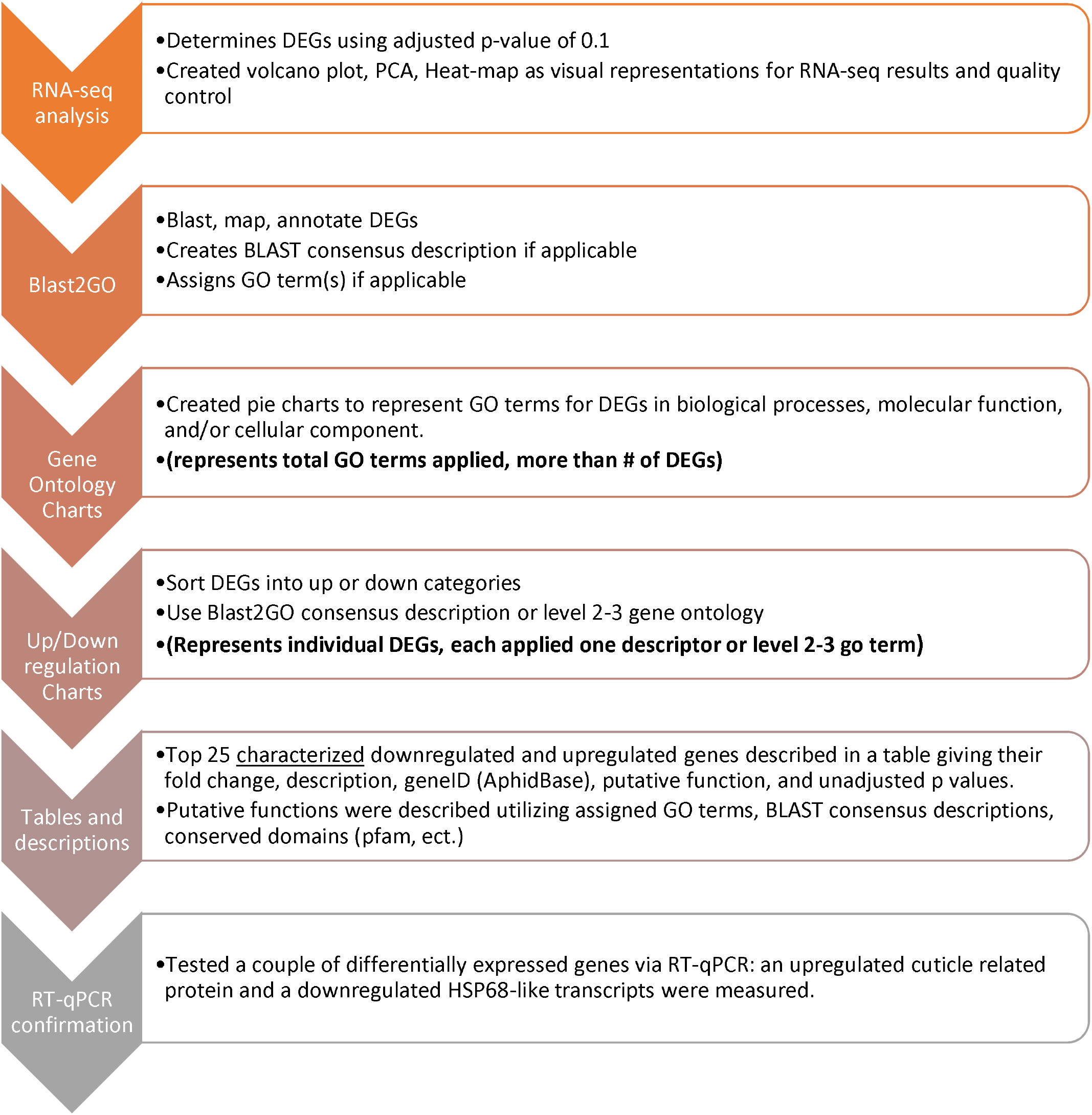
Explanation of process.

Supplementary Table S1: Primer Table

Supplementary Table S2: RNAseq significance list

Supplementary Table S3: Conserved σ32 −10 and −35 binding sites in *Escherichia coli* and *Buchnera* taxa

## References

1. Remaudiere G, Remaudiere M. Catalogue of the world’s Aphididae: Homoptera Aphidoidea. Paris, France: Institut National de la Recherche Agronomique (INRA); 1997. pp. 473–1275.

2. Fereres A, Irwin ME, Kamppeier GE. Aphid movement: process and consequences. In: van Emden HF, R. H, editors. Aphids as crop pests. 2nd ed. Oxfordshire, UK: CABI; 2017. p. 196–200.

3. Heck M. Insect Transmission of Plant Pathogens: a Systems Biology Perspective. mSystems®. 2018;3(2):e00168–17. doi: 10.1128/mSystems.00168-17.

4. Adams D, Douglas AE. How symbiotic bacteria influence plant utilisation by the polyphagous aphid, *Aphis fabae*. Oecologia. 1997;110(4):528–32. doi: 10.1007/s004420050190.

5. Pinheiro P V, Ghanim M, Alexander MM, Rebelo AR, Santos RS, Orsburn BC, et al. Host plants indirectly influence plant virus transmission by altering gut cysteine protease activity of aphid vectors. Mol Cell Proteomics. 2017;16(4 suppl 1):S230–43. doi: 10.1074/mcp.M116.063495.

6. Ng JCK, Perry KL. Transmission of plant viruses by aphid vectors. Mol Plant Pathol. 2004;5(5):505–11. doi: 10.1111/j.1364-3703.2004.00240.x.

7. Whitfield AE, Falk BW, Rotenberg D. Insect vector-mediated transmission of plant viruses. Virology. 2015;479–480:278–89. doi: 10.1016/j.virol.2015.03.026.

8. Elena SF, Bernet GP, Carrasco JL. The games plant viruses play. Curr Opin Virol. 2014;8:62–7. doi: 10.1016/j.coviro.2014.07.003.

9. Casteel CL, Falk BW. Plant virus-vector interactions: More than just for virus transmission. In: Wang A., Zhou X., editors. Current Research Topics in Plant Virology. 2016. p. 217 – 240. doi: 10.1007/978-3-319-32919-2_9.

10. Eigenbrode SD, Bosque-Pérez NA, Davis TS. Insect-borne plant pathogens and their vectors: ecology, evolution, and complex interactions. Annu Rev Entomol. 2018;63:169–191. doi: 10.1146/annurev-ento-020117-043119.

11. Blanc S, Michalakis Y. Manipulation of hosts and vectors by plant viruses and impact of the environment. Curr Opin Insect Sci. 2016; 16:36–43. doi: 10.1016/j.cois.2016.05.007.

12. Ingwell LL, Eigenbrode SD, Bosque-Pérez NA. Plant viruses alter insect behavior to enhance their spread. Sci Rep. 2012;2(1):578. doi: 10.1038/srep00578.

13. Stafford CA, Yang LH, Mcmunn MS, Ullman DE. Virus infection alters the predatory behavior of an omnivorous vector. Oikos. 2014;123:1384–1390. doi: 10.1111/oik.01148.

14. Wang Q, Li J, Dang C, Chang X, Fang Q, Stanley D, et al. *Rice dwarf virus* infection alters green rice leafhopper host preference and feeding behavior. PLoS One. 2018;13(9):1–16. doi: 10.1371/journal.pone.0203364.

15. Stafford CA, Walker GP, Ullman DE. Infection with a plant virus modifies vector feeding behavior. Proc Natl Acad Sci. 2011;108(23):9350–5. doi: 10.1073/pnas.1100773108.

16. Zhang YC, Cao WJ, Zhong LR, Godfray HCJ, Liu XD. Host plant determines the population size of an obligate symbiont (*Buchnera aphidicola*) in aphids. Appl Environ Microbiol. 2016;82(8):2336–46. doi: 10.1128/AEM.04131-15.

17. Hansen AK, Moran NA. Aphid genome expression reveals host-symbiont cooperation in the production of amino acids. Proc Natl Acad Sci. 2011;108(7):2849–54. doi: 10.1073/pnas.1013465108.

18. Nakabachi A, Shigenobu S, Sakazume N, Shiraki T, Hayashizaki Y, Carninci P, et al. Transcriptome analysis of the aphid bacteriocyte, the symbiotic host cell that harbors an endocellular mutualistic bacterium, *Buchnera*. Proc Natl Acad Sci. 2005;102(15):5477–82. doi: 10.1073/pnas.1013465108.

19. Wernegreen JJ. Strategies of genomic integration within insect-bacterial mutualisms. Biol Bull. 2012;223(1):112–22. doi: 10.1086/BBLv223n1p112.

20. Zhang Y, Su X, Harris A, Caraballo-Ortiz MA, Ren Z, Zhong Y. Genetic structure of the bacterial endosymbiont, *Buchnera aphidicola,* from its host aphid, *Schlechtendalia chinensis,* and evolutionary implications. Curr Microbiol. 2018;75(3):309–15. doi: 10.1007/s00284-017-1381-0.

21. Zhang F, Li X, Zhang Y, Coates B, Zhou XJ, Cheng D. Bacterial symbionts, *Buchnera*, and starvation on wing dimorphism in English grain aphid, *Sitobion avenae* (F.) (Homoptera: Aphididae). Front Physiol. 2015;6:155. doi: 10.3389/fphys.2015.00155.

22. Machado-Assefh CR, Lopez-Isasmendi G, Tjallingii WF, Jander G, Alvarez AE. Disrupting *Buchnera aphidicola*, the endosymbiotic bacteria of *Myzus persicae*, delays host plant acceptance. Arthropod Plant Interact. 2015;9(5):529–41. doi: 10.1007/s11829-015-9394-8.

23. Tamas I, Klasson L, Canbäck B, Näslund AK, Eriksson AS, Wernegreen JJ, et al. 50 Million years of genomic stasis in endosymbiotic bacteria. Science. 2002;296(5577):2376–9. doi: 10.1126/science.1071278.

24. Van Ham RCHJ, Kamerbeek J, Palacios C, Rausell C, Abascal F, Bastolla U, et al. Reductive genome evolution in *Buchnera aphidicola*. Proc Natl Acad Sci. 2003;100 (2) 581–586. doi: 10.1073/pnas.0235981100.

25. Douglas AE. Nutritional interactions in insect-microbial symbioses: aphids and their symbiotic bacteria *Buchnera*. Annu Rev Entomol. 1998;43(1):17–37. doi: 10.1146/annurev.ento.43.1.17.

26. Bouvaine S, Boonham N, Douglas AE. Interactions between a *Luteovirus* and the GroEL chaperonin protein of the symbiotic bacterium *Buchnera aphidicola* of aphids. J Gen Virol. 2011;92(6):1467–74. doi: 10.1099/vir.0.029355-0

27. Rana VS, Singh ST, Priya NG, Kumar J, Rajagopal R. *Arsenophonus* GroEL interacts with CLCuV and is localized in midgut and salivary gland of whitefly *B. tabaci*. PLoS One. 2012;7(8):e42168. doi: 10.1371/journal.pone.0042168.

28. Kliot A, Ghanim M. The role of bacterial chaperones in the circulative transmission of plant viruses by insect vectors. Viruses. 2013;5(6):1516–35. doi: 10.3390/v5061516.

29. Filichkin SA, Brumfield S, Filichkin TP, Young MJ. In vitro interactions of the aphid endosymbiotic SymL chaperonin with *Barley yellow dwarf virus*. J Virol. 1997;71(1):569–77. doi: 10.1128/JVI.71.1.569-577.1997.

30. van den Heuvel JF, Verbeek M, van der Wilk F. Endosymbiotic bacteria associated with circulative transmission of *Potato leafroll virus* by *Myzus persicae*. J Gen Virol. 1994;75 (Pt 10):2559–65. doi: 10.1099/0022-1317-75-10-2559.

31. Gray SM, Gildow FE. *Luteovirus*-aphid interactions. Annu Rev Phytopathol. 2003;41(1):539–66. doi: 10.1146/annurev.phyto.41.012203.105815

32. Li C, Cox-Foster D, Gray SM, Gildow F. Vector specificity of *Barley yellow dwarf virus* (BYDV) transmission: identification of potential cellular receptors binding BYDV-MAV in the aphid, *Sitobion avenae*. Virology. 2001;286(1):125–33. doi: 10.1006/viro.2001.0929.

33. Dombrovsky A, Gollop N, Chen S, Chejanovsky N, Raccah B. In vitro association between the helper component-proteinase of *Zucchini yellow mosaic virus* and cuticle proteins of *Myzus persicae*. J Gen Virol. 2007;88(5):1602–10. doi: 10.1099/vir.0.82769-0

34. van den Heuvel JF, Bruyère A, Hogenhout SA, Ziegler-Graff V, Brault V, Verbeek M, et al. The N-terminal region of the *luteovirus* readthrough domain determines virus binding to *Buchnera* GroEL and is essential for virus persistence in the aphid. Journal of Virology. 1997;71(10):7258–65. doi: 10.1128/JVI.71.10.7258-7265.1997.

35. Morin S, Ghanim M, Zeidan M, Czosnek H, Verbeek M, van den Heuvel JF. A GroEL homologue from endosymbiotic bacteria of the whitefly *Bemisia tabaci* is implicated in the circulative transmission of *Tomato yellow leaf curl virus*. Virology. 1999;256(1):75–84. doi: 10.1006/viro.1999.9631.

36. Chaudhary R, Atamian HS, Shen Z, Briggs SP, Kaloshian I. GroEL from the endosymbiont *Buchnera aphidicola* betrays the aphid by triggering plant defense. Proc Natl Acad Sci. 2014;111(24):8919–24. doi: 10.1073/pnas.1407687111.

37. Vandermoten S, Harmel N, Mazzucchelli G, De Pauw E, Haubruge E, Francis F. Comparative analyses of salivary proteins from three aphid species. Insect Mol Biol. 2014;23(1):67–77. doi: 10.1111/imb.12061.

38. Casteel CL, Hansen AK. Evaluating insect-microbiomes at the plant-insect interface. J Chem Ecol. 2014;40(7):836–47. doi: 10.1007/s10886-014-0475-4.

39. Gray SM, Cilia M, Ghanim M. Circulative, “nonpropagative” virus transmission: An orchestra of virus-, insect-, and plant-derived instruments. Adv Virus Res. 2014;89. doi: 10.1016/B978-0-12-800172-1.00004-5.

40. Eigenbrode SD, Ding H, Shiel P, Berger PH. Volatiles from potato plants infected with *Potato leafroll virus* attract and arrest the virus vector, *Myzus persicae* (Homoptera: Aphididae). Proc Biol Sci. 2002;269(1490):455–60. doi: 10.1098/rspb.2001.1909.

41. Heck M, Brault V. Targeted disruption of aphid transmission: a vision for the management of crop diseases caused by Luteoviridae members. Curr Opin Virol. 2018;33:24–32. doi: 10.1016/j.coviro.2018.07.007.

42. Rajabaskar D, Wu Y, Bosque-Pérez NA, Eigenbrode SD. Dynamics of *Myzus persicae* arrestment by volatiles from *Potato leafroll virus*-infected potato plants during disease progression. Entomol Exp Appl. 2013;148(2). doi: 10.1111/eea.12087.

43. Pinheiro P V, Wilson JR, Xu Y, Zheng Y, Rebelo AR, Fattah-Hosseini S, et al. Plant viruses transmitted in two different modes produce differing effects on small RNA-mediated processes in their aphid vector. Phytobiomes J. 2019;3(1):71. doi: 10.1094/PBIOMES-10-18-0045-R.

44. Hansen AK, Moran NA. Altered tRNA characteristics and 3′ maturation in bacterial symbionts with reduced genomes. Nucleic Acids Res. 2012;40(16):7870–84. doi: 10.1093/nar/gks503.

45. Thairu MW, Cheng S, Hansen AK. A sRNA in a reduced mutualistic symbiont genome regulates its own gene expression. Mol Ecol. 2018;27(8):1766–76. doi: 10.1111/mec.14424.

46. Thairu MW, Hansen AK. Changes in aphid host plant diet influence the small-RNA expression profiles of its obligate nutritional symbiont, *Buchnera*. mBio. 2019;10 (6) e01733–19. doi: 10.1128/mBio.01733-19.

47. Patton MF, Bak A, Sayre JM, Heck ML, Casteel CL. A polerovirus, *Potato leafroll virus*, alters plant–vector interactions using three viral proteins. Plant Cell Environ. 2020;43(2):387–399. doi: 10.1111/pce.13684.

48. Sadowy E, Juszczuk M, David C, Gronenborn B, Danuta Hulanicka MD. Mutational analysis of the proteinase function of *Potato leafroll virus*. J Gen Virol. 2001;82(Pt 6):1517–1527. doi: 10.1099/0022-1317-82-6-1517.

49. DeBlasio SL, Johnson R, Mahoney J, Karasev A, Gray SM, MacCoss MJ, et al. Insights into the *polerovirus*– plant interactome revealed by coimmunoprecipitation and mass spectrometry. Mol Plant-Microbe Interact. 2015;28(4):467–81. doi: 10.1094/MPMI-11-14-0363-R.

50. Zhong S, Joung JG, Zheng Y, Chen YR, Liu B, Shao Y, et al. High-throughput illumina strand-specific RNA sequencing library preparation. Cold Spring Harb Protoc. 2011;2011(8):940–9. doi: 10.1101/pdb.prot5652.

51. Anders S, McCarthy DJ, Chen Y, Okoniewski M, Smyth GK, Huber W, et al. Count-based differential expression analysis of RNA sequencing data using R and Bioconductor. Nat Protoc. 2013;8(9):1765–86. doi: 10.1038/nprot.2013.099.

52. Morgan M, Anders S, Lawrence M, Aboyoun P, Pagès H, Gentleman R. ShortRead: A bioconductor package for input, quality assessment and exploration of high-throughput sequence data. Bioinformatics. 2009;25(19):2607–8. doi: 10.1093/bioinformatics/btp450.

53. Andrews S. FastQC: A quality control tool for high throughput sequence data. http://www.bioinformatics.babraham.ac.uk/projects/fastqc/. 2010.

54. Gauthier JP, Legeai F, Zasadzinski A, Rispe C, Tagu D. AphidBase: A database for aphid genomic resources. Bioinformatics. 2007;23(6):783–4. doi: 10.1093/bioinformatics/btl682.

55. Kim D, Pertea G, Trapnell C, Pimentel H, Kelley R, Salzberg SL. TopHat2: Accurate alignment of transcriptomes in the presence of insertions, deletions and gene fusions. Genome Biol. 2013;14:R36. doi: 10.1186/gb-2013-14-4-r36.

56. Anders S, Pyl PT, Huber W. HTSeq-A Python framework to work with high-throughput sequencing data. Bioinformatics. 2015;31(2):166–9. doi: 10.1093/bioinformatics/btu638.

57. Love MI, Huber W, Anders S. Moderated estimation of fold change and dispersion for RNA-seq data with DESeq2. Genome Biol. 2014;15(12):550. doi: 10.1186/s13059-014-0550-8.

58. Conesa A, Götz S, García-Gómez JM, Terol J, Talón M, Robles M. Blast2GO: A universal tool for annotation, visualization and analysis in functional genomics research. Bioinformatics. 2005;21(18):3674–6. doi: 10.1093/bioinformatics/bti610

59. Patton MF, Arena GD, Salminen JP, Steinbauer MJ, Casteel CL. Transcriptome and defence response in *Eucalyptus camaldulensis* leaves to feeding by *Glycaspis brimblecombei* Moore (Hemiptera: Aphalaridae): a stealthy psyllid does not go unnoticed. Austral Entomol. 2017;57(2):247–54. doi: 10.1111/aen.12319

60. Casteel CL, De Alwis M, Bak A, Dong H, Whitham SA, Jander G. Disruption of ethylene responses by *Turnip mosaic virus* mediates suppression of plant defense against the green peach aphid vector. Plant Physiol. 2015;169(1):209–18. doi: 10.1104/pp.15.00332

61. Nikoh N, McCutcheon JP, Kudo T, Miyagishima SY, Moran NA, Nakabachi A. Bacterial genes in the aphid genome: Absence of functional gene transfer from Buchnera to its host. PLoS Genet. 2010;6(2):e1000827. doi: 10.1371/journal.pgen.1000827.

62. Hansen AK, Degnan PH. Widespread expression of conserved small RNAs in small symbiont genomes. ISME J. 2014;8(12):2490–502. doi: 10.1038/ismej.2014.121.

63. Hogenhout SA, van der Wilk F, Verbeek M, Goldbach RW, van den Heuvel JF. *Potato leafroll virus* binds to the equatorial domain of the aphid endosymbiotic GroEL homolog. J Virol. 1998;72(1):358–65. doi: 10.1128/JVI.72.1.358-365.1998.

64. Camberg JL, Doyle SM, Johnston DM, Wickner S. Molecular Chaperones. In: Brenner’s Encyclopedia of Genetics: Second Edition. Elsevier; 2013. p. 456–60. doi: 10.1016/B978-0-12-809633-8.06723-6.

65. Segal G, Ron EZ. Regulation and organization of the *groE* and *dnaK* operons in Eubacteria. FEMS Microbiol Lett. 1996;138(1):1–10. doi: 10.1111/j.1574-6968.1996.tb08126.x

66. Zhang L, Pelech S, Uitto VJ. Bacterial GroEL-like heat shock protein 60 protects epithelial cells from stress-induced death through activation of ERK and inhibition of caspase 3. Exp Cell Res. 2004;292(1):231–40. doi: 10.1016/j.yexcr.2003.08.012.

67. Dombrovsky A, Sobolev I, Chejanovsky N, Raccah B. Characterization of RR-1 and RR-2 cuticular proteins from *Myzus persicae*. Comp Biochem Physiol - B Biochem Mol Biol. 2007;146(2):256–64. doi: 10.1016/j.cbpb.2006.11.013.

68. Dombrovsky A, Huet H, Zhang H, Chejanovsky N, Raccah B. Comparison of newly isolated cuticular protein genes from six aphid species. Insect Biochem Mol Biol. 2003;33(7):709–15. doi: 10.1016/s0965-1748(03)00065-1.

69. Liang Y, Gao XW. The cuticle protein gene MPCP4 of *Myzus persicae* (Homoptera: Aphididae) plays a critical role in cucumber mosaic virus acquisition. J Econ Entomol. 2017;110(3):848–53. doi: 10.1093/jee/tox025

70. Silva AX, Jander G, Samaniego H, Ramsey JS, Figueroa CC. Insecticide resistance mechanisms in the green peach aphid *Myzus persicae* (Hemiptera: Aphididae) I: A transcriptomic survey. PLoS One. 2012;7(6):e36366. doi: 10.1371/journal.pone.0036366

71. Deshoux M, Monsion B, Uzest M. Insect cuticular proteins and their role in transmission of phytoviruses. Curr Opin Virol. 2018;33:137–143. doi: 10.1016/j.coviro.2018.07.015.

72. Gallot A, Rispe C, Leterme N, Gauthier JP, Jaubert-Possamai S, Tagu D. Cuticular proteins and seasonal photoperiodism in aphids. Insect Biochem Mol Biol. 2010;40(3):235–40. doi: 10.1016/j.ibmb.2009.12.001.

73. Cilia M, Tamborindeguy C, Fish T, Howe K, Thannhauser TW, Gray SM. Genetics coupled to quantitative intact proteomics links heritable aphid and endosymbiont protein expression to circulative polerovirus transmission. J Virol. 2011;85(5):2148–66. doi: 10.1128/JVI.01504-10.

74. Wang H, Wu K, Liu Y, Wu Y, Wang X. Integrative proteomics to understand the transmission mechanism of *Barley yellow dwarf virus*-GPV by its insect vector *Rhopalosiphum padi*. Sci Rep. 2015;5:10971. doi: 10.1038/srep10971.

75. Seddas P, Boissinot S, Strub JM, Van Dorsselaer A, Van Regenmortel MHV, Pattus F. Rack-1, GAPDH3, and actin: proteins of *Myzus persicae* potentially involved in the transcytosis of *Beet western yellows virus* particles in the aphid. Virology. 2004;325(2):399–412. doi: 10.1016/j.virol.2004.05.014

76. Yang Z, Zhang F, Zhu L, He G. Identification of differentially expressed genes in brown planthopper *Nilaparvata lugens* (Hemiptera: Delphacidae) responding to host plant resistance. Bull Entomol Res. 2006;96(1):53–9. doi: 10.1079/ber2005400.

77. Bass C, Zimmer CT, Riveron JM, Wilding CS, Wondji CS, Kaussmann M, et al. Gene amplification and microsatellite polymorphism underlie a recent insect host shift. Proc Natl Acad Sci. 2013;110(48):19460–5. doi: 10.1073/pnas.1314122110

78. Ramsey JS, Elzinga DA, Sarkar P, Xin YR, Ghanim M, Jander G. Adaptation to nicotine feeding in *Myzus persicae*. J Chem Ecol. 2014;40(8):869–77. doi: 10.1007/s10886-014-0482-5

79. Casteel CL, Jander G. New synthesis: investigating mutualisms in virus-vector interactions. J Chem Ecol. 2013;39(7):809. doi: 10.1007/s10886-013-0305-0.

80. Götz M, Popovski S, Kollenberg M, Gorovits R, Brown JK, Cicero JM, et al. Implication of *Bemisia tabaci HEAT SHOCK PROTEIN* 70 in *Begomovirus*-whitefly interactions. J Virol. 2012;86(24):13241–52. doi: 10.1128/JVI.00880-12.

81. Porras MF, Navas CA, Marden JH, Mescher MC, De Moraes CM, Pincebourde S, et al. Enhanced heat tolerance of viral-infected aphids leads to niche expansion and reduced interspecific competition. Nat Commun. 2020;11(1):1184. doi: 10.1038/s41467-020-14953-2.

82. Syller J. The influence of temperature on transmission of potato leaf roll virus by *Myzus persicae* Sulz. Potato Res. 1987;30(1):47–58. doi: 10.1007/BF02357683.

83. Syller J. The Effects of Temperature on the Susceptibility of Potato Plants to Infection and Accumulation of *Potato Leafroll Virus*. J Phytopathol. 1991;133(3):216–24. doi: 10.1111/j.1439-0434.1991.tb00156.x

84. Chung BN, Canto T, Tenllado F, Choi KS, Joa JH, Ahn JJ, et al. The effects of high temperature on infection by *Potato virus Y*, *Potato virus A*, and *Potato leafroll virus*. Plant Pathol J. 2016;32(4):321–8. doi: 10.5423/PPJ.OA.12.2015.0259.

85. Mandrioli M, Manicardi GC. Chromosomal mapping reveals a dynamic organization of the histone genes in aphids (Hemiptera: Aphididae). Entomologia. 2013 1(1), e2. doi: 10.4081/entomologia.2013.e2.

86. Enders LS, Bickel RD, Brisson JA, Heng-Moss TM, Siegfried BD, Zera AJ, et al. Abiotic and biotic stressors causing equivalent mortality induce highly variable transcriptional responses in the soybean aphid. G3 (Bethesda). 2014;5(2):261–70. doi: 10.1534/g3.114.015149.

87. Prado F, Jimeno-Gonz alez S, Reyes JC. Histone availability as a strategy to control gene expression. RNA Biol. 2017;14(3):281–286. doi: 10.1080/15476286.2016.1189071.

88. Shigenobu S, Watanabe H, Hattori M, Sakaki Y, Ishikawa H. Genome sequence of the endocellular bacterial symbiont of aphids *Buchnera* sp. APS. Nature. 2000;407(6800):81–6. doi: 10.1038/35024074.

89. Jiang Z, Jones DH, Khuri S, Tsinoremas NF, Wyss T, Jander G, et al. Comparative analysis of genome sequences from four strains of the *Buchnera aphidicola* Mp endosymbion of the green peach aphid, *Myzus persicae*. BMC Genomics. 2013;14(1):917. doi: 10.1186/1471-2164-14-917

90. Wilcox JL, Dunbar HE, Wolfinger RD, Moran NA. Consequences of reductive evolution for gene expression in an obligate endosymbiont. Mol Microbiol. 2003;48(6):1491–500. doi: 10.1046/j.1365-2958.2003.03522.x.

91. Karp PD, Billington R, Caspi R, Fulcher CA, Latendresse M, Kothari A, et al. The BioCyc collection of microbial genomes and metabolic pathways. Brief Bioinform. 2019;20(4):1085–1093. doi: 10.1093/bib/bbx085.

92. Gross CA. Function and regulation of the heat shock proteins. 2nd ed. Neidhardt FC, editor. Vol. 1, Escherichia coli and Salmonella: Cellular and Molecular Biology. Washington DC: American Society for Microbiology Press; 1996. 1382–1399 p.

93. Chong RA, Moran NA. Intraspecific genetic variation in hosts affects regulation of obligate heritable symbionts. Proc Natl Acad Sci. 2016;113(46):13114–9. doi: 10.1073/pnas.1610749113.

94. Pers D, Hansen AK. The boom and bust of the aphid’s essential amino acid metabolism across nymphal development. G3 (Bethesda). 2021:jkab115. doi: 10.1093/g3journal/jkab115.

95. Zhang B, Leonard SP, Li Y, Moran NA. Obligate bacterial endosymbionts limit thermal tolerance of insect host species. Proc Natl Acad Sci. 2019;116(49):24712–8. doi: 10.1073/pnas.1915307116.

96. Dunbar HE, Wilson ACC, Ferguson NR, Moran NA. Aphid thermal tolerance is governed by a point mutation in bacterial symbionts. PLoS Biol. 2007;5(5):e96. doi: 10.1371/journal.pbio.0050096.

97. Moran NA, Yun Y. Experimental replacement of an obligate insect symbiont. Proc Natl Acad Sci. 2015;112(7):2093–6. doi: 10.1073/pnas.1420037112

98. Fares MA, Barrio E, Sabater-Muñoz B, Moya A. The evolution of the heat-shock protein GroEL from *Buchnera*, the primary endosymbiont of aphids, is governed by positive selection. Mol Biol Evol. 2002;19(7):1162–70. doi: 0.1093/oxfordjournals.molbev.a004174

99. Kliot A, Cilia M, Czosnek H, Ghanim M. Implication of the bacterial endosymbiont *Rickettsia* spp. in interactions of the whitefly *Bemisia tabaci* with *Tomato yellow leaf curl virus*. J Virol. 2014; 15;88(10):5652–60. doi: 10.1128/JVI.00071-14

100. Hosseinzadeh S, Shams-Bakhsh M, Mann M, Fattah-Hosseini S, Bagheri A, Mehrabadi M, et al. Distribution and variation of bacterial endosymbiont and “*Candidatus* Liberibacter asiaticus” titer in the Huanglongbing insect vector, *Diaphorina citri Kuwayama*. Microb Ecol. 2019;78(1):206–222. doi: 10.1007/s00248-018-1290-1.

101. Ramsey JS, Johnson RS, Hoki JS, Kruse A, Mahoney J, Hilf ME, et al. Metabolic interplay between the asian citrus psyllid and its profftella symbiont: An achilles’ heel of the citrus greening insect vector. PLoS One. 2015;10(11):e0140826. doi: 10.1371/journal.pone.0140826.

102. Smith TE, Moran NA. Coordination of host and symbiont gene expression reveals a metabolic tug-of-war between aphids and Buchnera. Proc Natl Acad Sci. 2020;117(4):2113–21. doi: 10.1073/pnas.1916748117

